# napari-threedee: a toolkit for human-in-the-loop 3D image analysis in napari

**DOI:** 10.1101/2023.07.28.550950

**Authors:** Kevin A. Yamauchi, Alister Burt

## Abstract

With recent advances in three-dimensional bioimaging acquisition and analysis methods, imaging data are an important tool for unveiling the complex architecture of biological organisms. However, extracting insights from these complex datasets requires interactive exploration as well as annotation for training supervised analytical methods. Here we present napari-threedee, a library of composable components and napari plugins for interactively exploring and annotating image data in 3D. We demonstrate how napari-threedee can be used for human-in-the-loop 3D image analysis in diverse image data including fluorescence microscopy and cryoelectron tomography.

---

Owing to improvements in imaging technology and computational methods, there are increasingly complex three dimensional imaging datasets being acquired. These improvements have led to biological insights at length scales ranging from molecular to whole orgaism length scales^1,2^. Fortunately, there has been significant improvements in automated image processing and a rise in viewers for visualizing three dimensional imaging data and analytical methods for extracting useful features. napari is a GPU-accelerated, python-based multidimensional image viewer that provides rendering for 2D and 3D datasets. However, these visualizations are largely static and do not allow users to easily modify and interact with their data in 3D (Supplementary Note 1). Further, many cutting edge computer vision require annotated ground truth annotations for training and/or benefit from human-in-the-loop supervision^3^. Thus, there is a need for software that enables users and developers to create rich, interactive analysis tools in 3D. Here, we report napari-threedee, a library for interactive annotation, exploration, and visualization of 3D data comprising a toolkit of readymade plugins for 3D interaction for napari^4^ users and composable components to enhance napari plugins with 3D interactivity. We demonstrate the broad applicability of napari-threedee with example workflows for particle picking for cryoelectron tomography and interactive, GPU-accelerated 3D segmentation of fluorescence microscopy data.

napari-threedee is a toolkit for 3D interactivity in napari that enables human in the loop analysis for 3D data (Figure 1A). napari-threedee provides interactivity for two types of users: (1) analysts and (2) developers. For the analysts, napari-threedee has napari plugins that bring the main types of interaction to native napari visualizations (Figure 1C). For developers, napari-threedee provides composable widgets that can be integrated into existing plugins to enable 3D interactivity. By targeting napari, napari-threedee enhanced workflows can be integrated with the scientific python ecosystem including numpy, scipy, pytorch, and tensorflow. Importantly, we have designed napari-threedee such that the objects can be interacted with both through the graphical user interface and via the python API. The state of the object can be then read back into the analysis pipeline, thus, closing the loop.

**Fig. 1.**
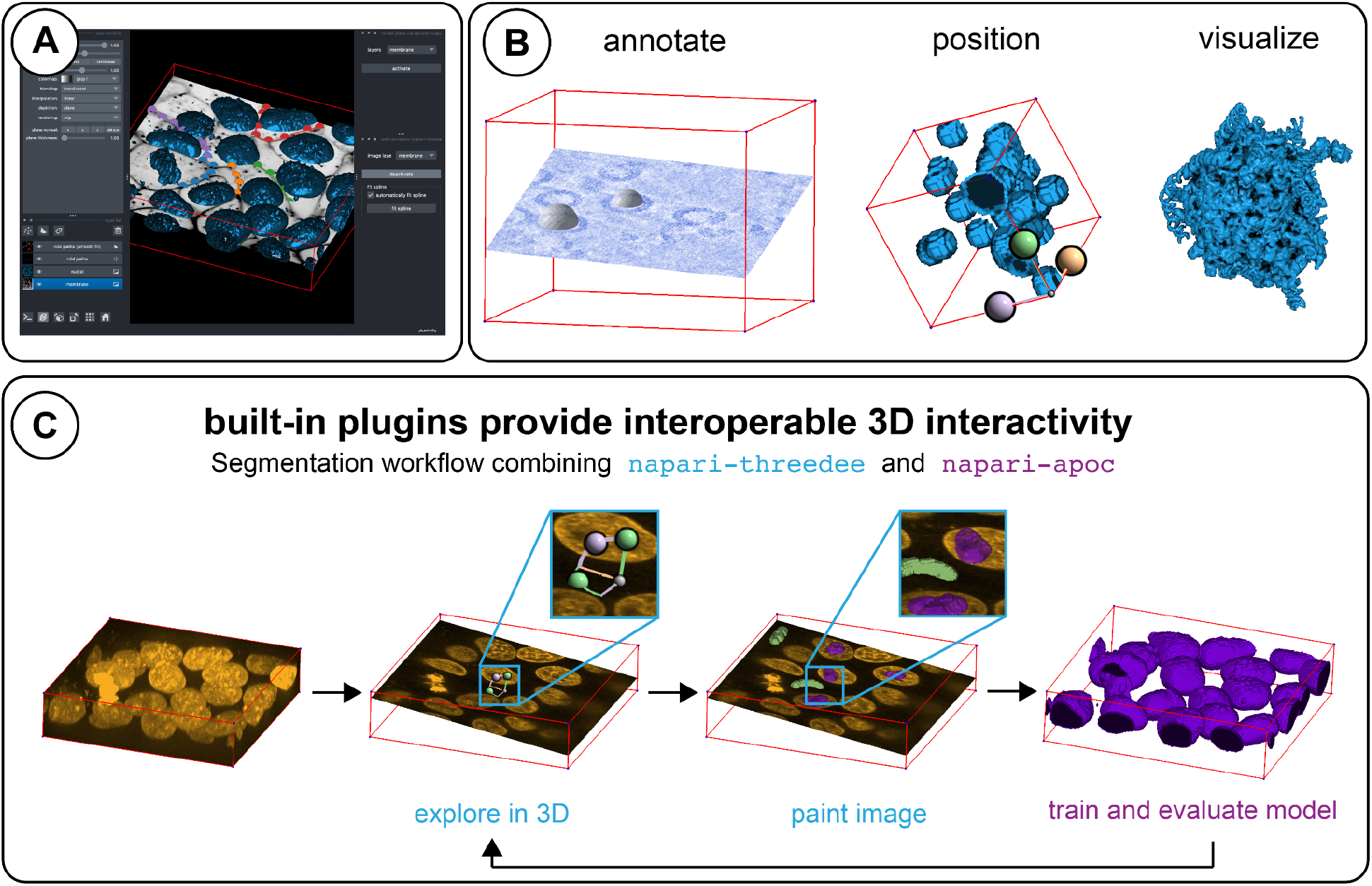
napari-threedee is a modular toolking for 3D human in the loop image processing in napari. **(a)** napari-threedee enables rich interactive human-in-the-loop analysis of 3D imaging data. **(b)** napari-threedee provides graphical user interfaces for interactive annotation and positioning of 3D data. Additionally, to improve napari-threedee has components to augment the visualization of 3D data in napari. **(c)** napari-threedee plugins allow for interaction out of the box and are interoperable with other plugins in the napari plugin ecosystem. Here we show how napari-threedee can be combined with napari-apoc for interactive 3D segmentation of fluorescently-labeled nuclei.

To interact with objects in a 3D dimensional scene, one must determine which visual is being clicked on and map the mouse movements to 3D movements. Mapping the 2D gestures on the screen to the 3D scene is non-trivial. We have implemented infrastructure for selecting and manipulating objects in a 3D scene in napari (Supplementary Note 2). To make this fundamental tooling available to as many users as possible, we have contributed much of it back to the upstream napari and vispy libraries (Supplementary Table 2).

Annotators provide GUI elements for interactive annotation in 3D. Annotation is an essential step in image analysis pipelines for many different workflows such as setting landmarks for registration or labeling the data for training machine learning algorithms. Annotators have enabled applications such as filament picking in cryoEM and editing of dense labels in 3D for cell segmentation in fluorescence microscopy. See Supplementary Note 3 for a description of the different types of annotators.

Manipulators provide GUI elements for interactively positioning objects in the scene in 3D. Positioning is useful for specifying the location or orientation of particles in CryoEM or a rendering plane for intuitively viewing an oblique slice in 3D. napari-threedee provides manipulators for positioning points, image layers, and rendering planes (Supplementary Note 4).

Visualizers augment the 3D data visualization capabilities of napari. We have a component for controlling the lighting of surface layer meshes. Additionally, we have created a component for applying ambient occlusion to meshes to add shadows to meshes. Together the visualizers improve the rendering of meshes in napari (Supplementary Note 5).

For analysts who wish to add 3D interactivity to their existing workflow, we have implemented 11 built-in plugins. By integrating with standard napari layer types, the napari-threedee plugins are interoperable with other napari plugins. As an example of how napari-threedee can be used with other plugins, we show how manipulators and annotators can be used with napari-apoc^5^ to create a GPU-accelerated 3D segmentation workflow (Figure 1C).

For developers who wish to add 3D interactivity to their plugins, we have provided composable components. The components have a model which controls the state of the interactive component and a visual that is placed in the rendered scene. There is bidirectional synchronization between the model and visual, which means that the visual can be positioned either via the GUI or the command line and the state of the visual can be used in the analysis. The model-view separation simplifies the integration of n3d components with existing workflows and applications. napari-tomoslice, a particle picking plugin for cryoelectron tomography is an example of how napari-threedee components can be used to create plugins with rich 3D interactivity.

To help users learn to use napari-threedee, we have provided documentation with examples, explanations, and tutorials (Supplementary Note 6). The provided examples show analysts how to use the provided plugins to add 3D interactivity to their existing workflows. The explanations are an educational resource that teaches users and developers how napari-threedee works. The tutorials give developers step-by-step instructions on how to integrate napari-threedee components into their plugins.

napari-threedee is a toolkit of readymade plugins and composable components for rich 3D interactivity in napari analysis workflows. napari-threedee addresses the need for 3D interactivity for both analysts and developers. We anticipate that users across the imaging community will be able to integrate 3D interactivity into their workflows.

## Code availability

napari-threedee is free open source software (BSD-3 license) and is available at: https://github.com/napari-threedee/napari-threedee. Documentation is available at: https://napari-threedee.github.io.

## Supplementary Material

### Supplementary Note 1: Landscape of 3D interactivity in image viewers

#### The need for 3D interactivity

Three dimensional imaging is an important tool for understanding the organization and structure of the natural world. Recent advances in imaging technology has resulted in a significant increase in throughput. In order to generate hypotheses from image data, it is often helpful for researchers to be able to easily explore and interact with their data. Doing so in 3D rather than 2D gives the researcher richer context for their observations. Extracting insights from large image datasets requires automated image processing. Advances in deep learning have greatly improved the speed and accuracy of many common computer vision tasks such as segmentation or classification. Many image analysis tasks require annotation

#### 3D interactivity in current image viewers

There are several open source image viewers with vibrant communities such as FIJI, ImageJ, and napari. These viewers all offer interactive annotation of images in 2D (e.g., draw regions of interest, annotate points) and can be extended via plugins, but they do not support interactive 3D annotation (Supplementary Table 1). On the other hand, Imaris does offer the ability to annotate in 3D, but is not open source. Given that no open source image viewer offers 3D interactivity, there is a need to extend a viewer with the capability.

**Supplementary Table 2:** Current open source image viewers lack 3D interactivity.

| Viewer | 2D interactivity | 3D interactivity | Open source |
| --- | --- | --- | --- |
| FIJI | Yes | No | Yes |
| ImageJ | Yes | No | Yes |
| Imaris | Yes | Yes | No |
| napari | Yes | No | Yes |
| napari +<br>napari-threedee | Yes | Yes | Yes |

### Supplementary Note 2: Architecture for 3D interactivity infrastructure

**Note:** we have contributed this explanation upstream to the core napari documentation. We will maintain the documentation there and recommend viewing it for the most up to date explanation. The documentation is here: https://napari.org/stable/guides/3D_interactivity.html

#### Coordinate systems in napari

In napari, there are three main coordinate systems: (1) canvas, (2) world, and (3) layer. The canvas coordinates system is the 2D coordinate system of the canvas on which the scene is rendered. World coordinates are the nD coordinates of the entire scene. As the name suggests, layer coordinates are the nD coordinate system of the data in a given layer. Layer coordinates are specific to each layer’s data and are related to the world coordinate system via the layer transforms.

**Supplementary Fig. 1.**
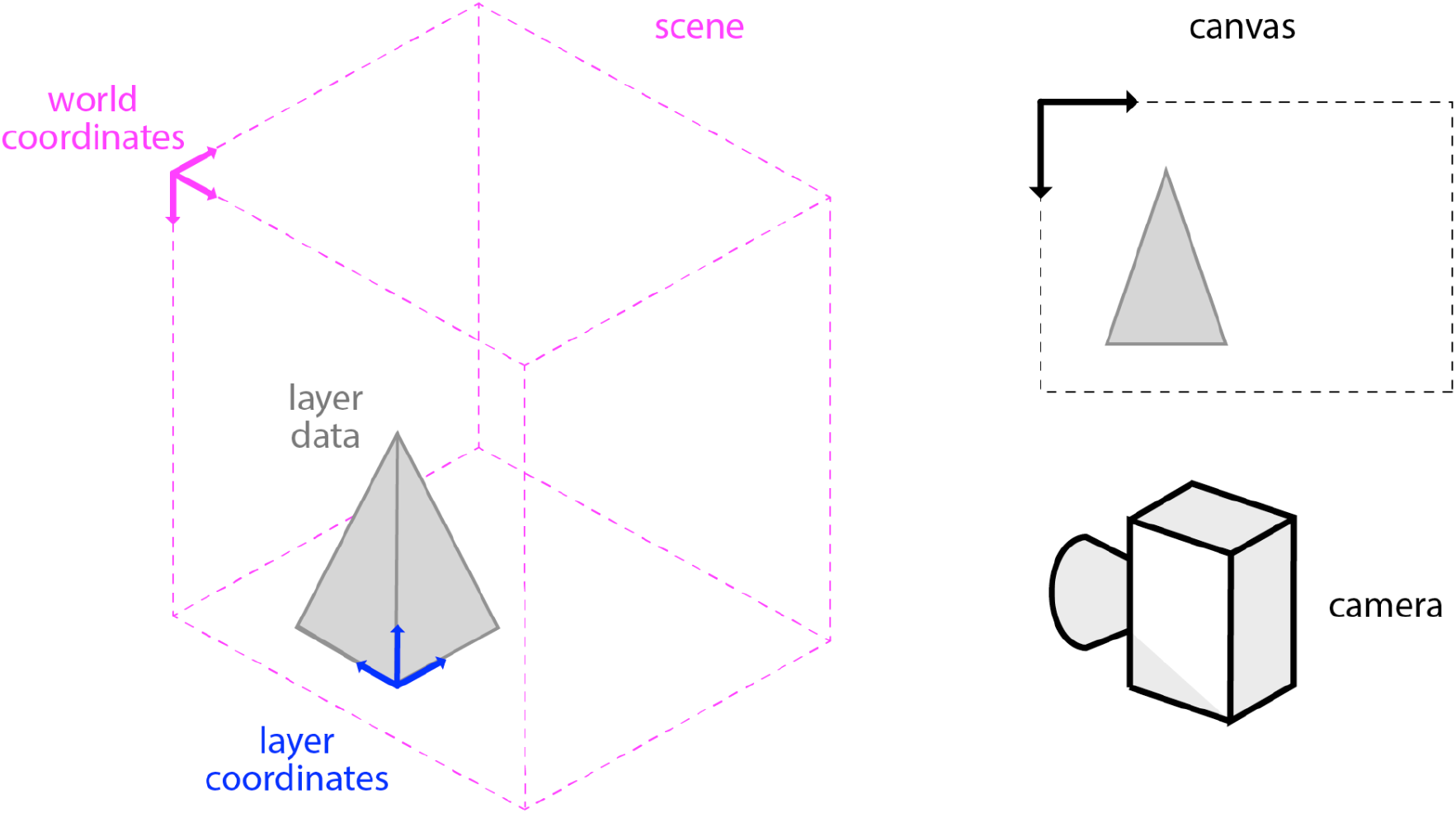
Overview of coordinate systems in napari. napari has three coordinate systems that must be transformed between in order to determine where mouse clicks correspond to in a given layer’s data.

#### In 3D mode, clicks are lines

Since the 3D scene is rendered on a 2D surface (your screen), your mouse click does not map to a specific point in space. As the view is a parallel projection, napari can determine a line through 3D space that intersects the canvas where the user clicked.

**Supplementary Fig. 2.**
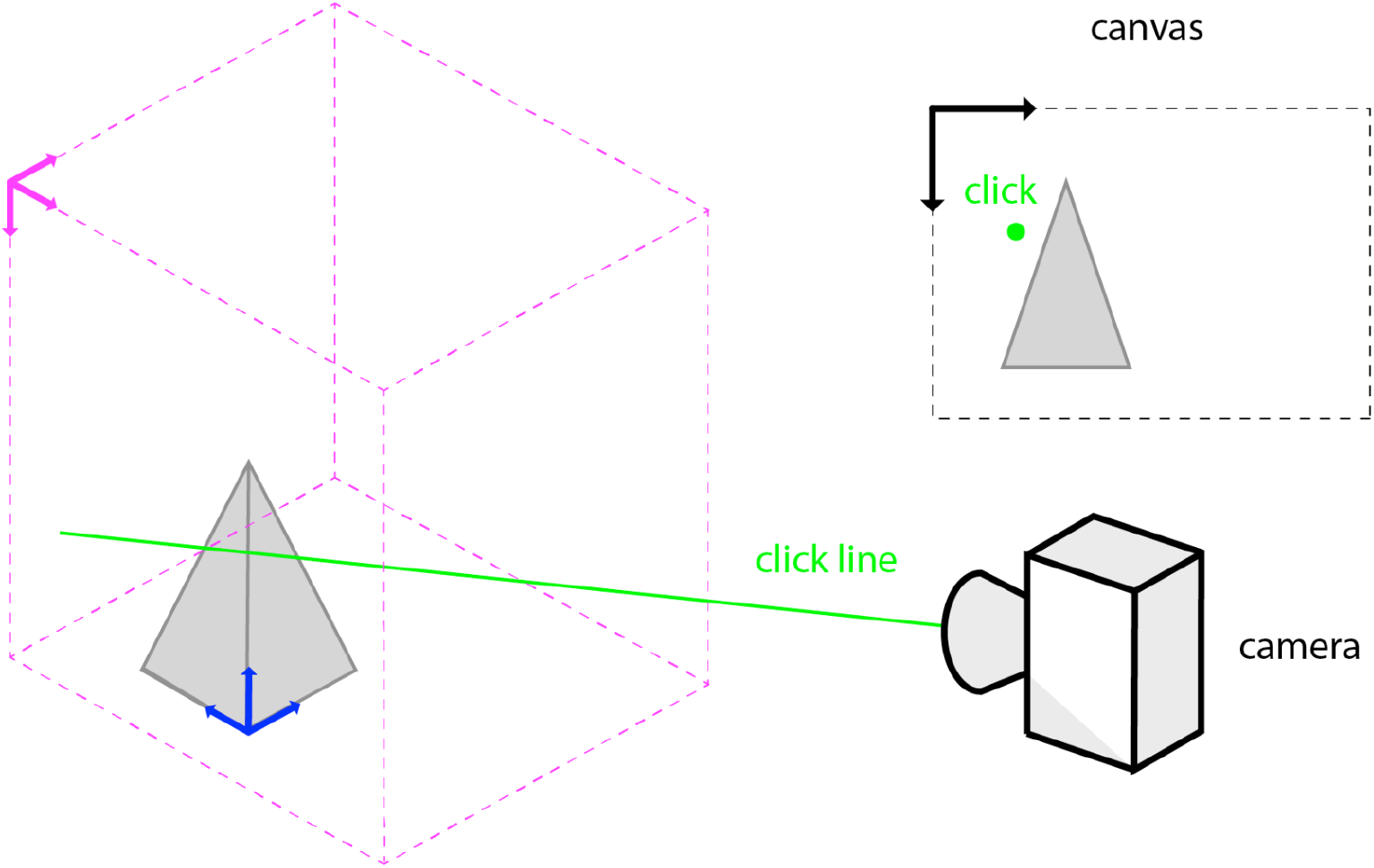
In a napari three dimensional scene, a click is projected as a line through the scene.

#### Determining where the click intersects the data

Each napari layer has a method called `get_ray_intersections()` that will return the points on the data bounding box that a given line will intersect (`start_point` and `end_point`). When the click line (`view_direction`) and position (`position`) are used as inputs, `start_point` and `end_point` are the end points of the segment click line that intersects the layer’s axis-alinged data bounding box. `start_point` is the end point that is closest to the camera (i.e, the “first” intersection) and `end_point` is the end point that is farthest from the camera (i.e., the “last” intersection). You can use the line segment between `start_point` and `end_point` to interrogate the layer data that is “under” your cursor.

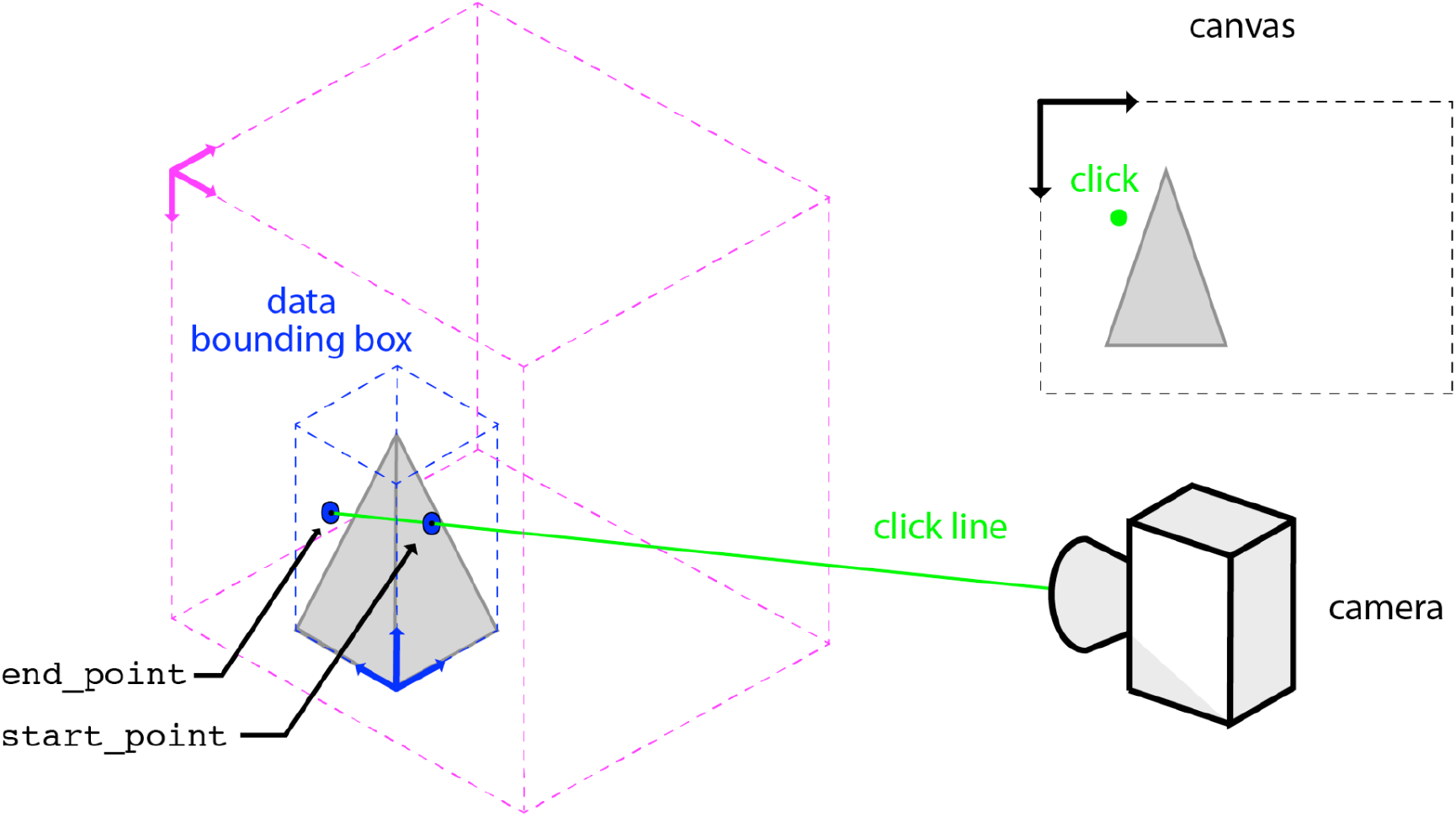

#### Getting the layer data under the cursor

There are convenience methods in the layer objects (`layer.get_value()`) to get the layer data value underneath the cursor that is “on top” (i.e., closest to `start_point`). Like `layer.get_ray_intersections()`, `layer.get_value()` takes the click position, view direction, dims_displayed in either world or layer coordinates (see `world` argument) as input. Thus, it can be easily integrated into a mouse event callback. Note that `layer.get_value()` returns `None` if the layer is not currently visible. See the docstring below for details.

**Supplementary Table 2:**
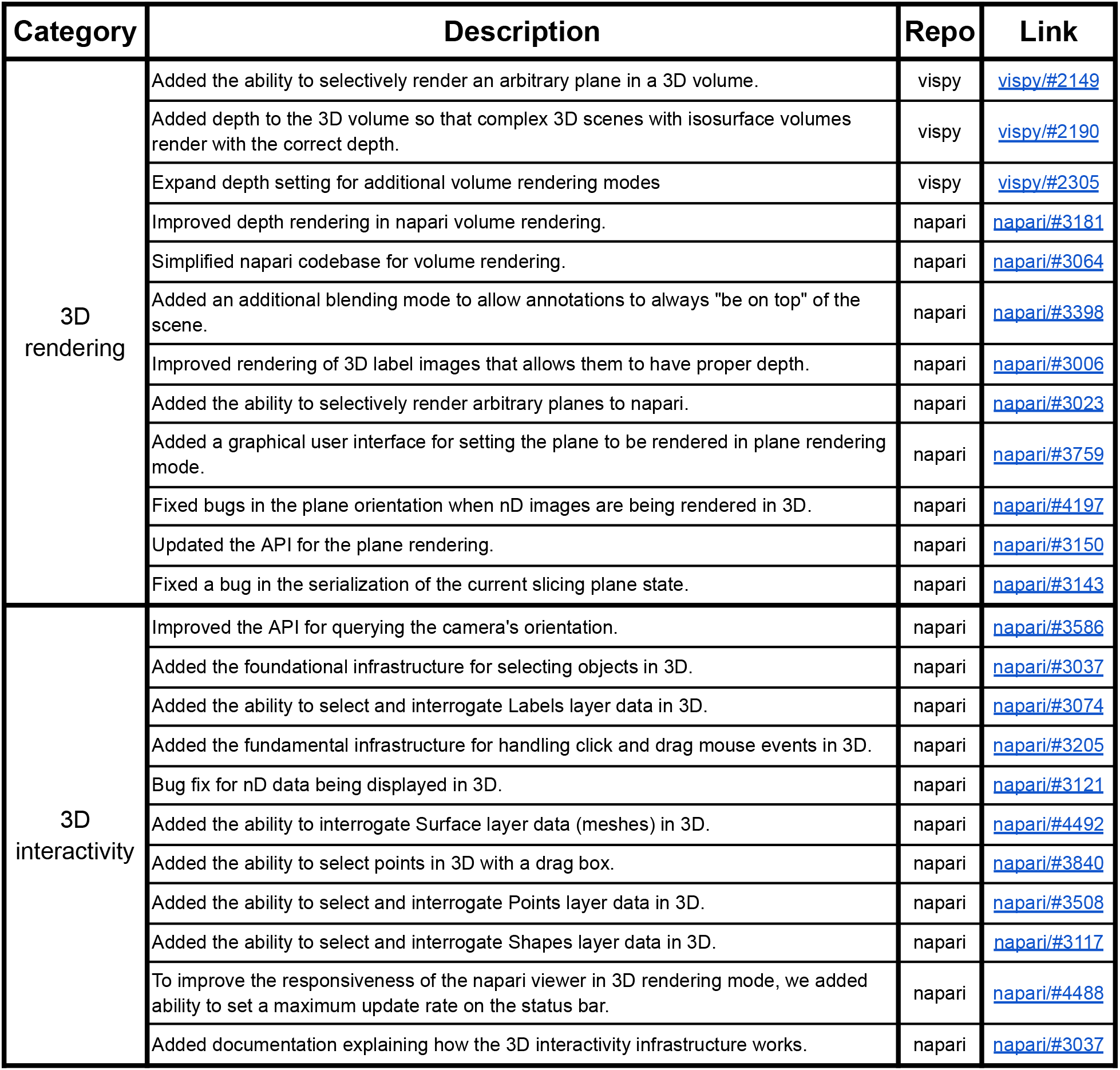
To make the improvements we made to enable 3D interactivity available to the broadest community, we contributed many of them to the upstream libraries we depend on for rendering and interactivity. This table lists contributions we made to improve 3D rendering and 3D interactivity in napari and vispy.

| Category | Description | Repo | Link |
| --- | --- | --- | --- |
| 3D rendering | Added the ability to selectively render an arbitrary plane in a 3D volume. | vispy | vispy/#2149 |
|  | Added depth to the 3D volume so that complex 3D scenes with isosurface volumes render with the correct depth. | vispy | vispy/#2190 |
|  | Expand depth setting for additional volume rendering modes | vispy | vispy/#2305 |
|  | Improved depth rendering in napari volume rendering. | napari | napari/#3181 |
|  | Simplified napari codebase for volume rendering. | napari | napari/#3064 |
|  | Added an additional blending mode to allow annotations to always "be on top" of the scene. | napari | napari/#3398 |
|  | Improved rendering of 3D label images that allows them to have proper depth. | napari | napari/#3006 |
|  | Added the ability to selectively render arbitrary planes to napari. | napari | napari/#3023 |
|  | Added a graphical user interface for setting the plane to be rendered in plane rendering mode. | napari | napari/#3759 |
|  | Fixed bugs in the plane orientation when nD images are being rendered in 3D. | napari | napari/#4197 |
|  | Updated the API for the plane rendering. | napari | napari/#3150 |
|  | Fixed a bug in the serialization of the current slicing plane state. | napari | napari/#3143 |
| 3D interactivity | Improved the API for querying the camera's orientation. | napari | napari/#3586 |
|  | Added the foundational infrastructure for selecting objects in 3D. | napari | napari/#3037 |
|  | Added the ability to select and interrogate Labels layer data in 3D. | napari | napari/#3074 |
|  | Added the fundamental infrastructure for handling click and drag mouse events in 3D. | napari | napari/#3205 |
|  | Bug fix for nD data being displayed in 3D. | napari | napari/#3121 |
|  | Added the ability to interrogate Surface layer data (meshes) in 3D. | napari | napari/#4492 |
|  | Added the ability to select points in 3D with a drag box. | napari | napari/#3840 |
|  | Added the ability to select and interrogate Points layer data in 3D. | napari | napari/#3508 |
|  | Added the ability to select and interrogate Shapes layer data in 3D. | napari | napari/#3117 |
|  | To improve the responsiveness of the napari viewer in 3D rendering mode, we added ability to set a maximum update rate on the status bar. | napari | napari/#4488 |
|  | Added documentation explaining how the 3D interactivity infrastructure works. | napari | napari/#3037 |

### Supplementary Note 3: Annotators

napari-threedee annotators provide methods to directly annotate data in 3D. Annotating data in 3D allows the person making the annotations to view the full context. This is important because many annotations span all three dimensions (e.g., filaments in cryoelectron tomography data or cells in 3D fluorescence microscopy images) and thus it can be difficult to annotate in axis-aligned 2D slices. We have developed 5 different types of annotators:

- **Plane labeler:** paint labels (e.g., segmentation ground truth) on arbitrary slicing planes in 3D.
- **PathAnnotator**: draw paths through the data defined by 3D splines. This is useful for labeling filaments, defining camera paths for exploring the data, and labeling anatomical features.
- **PointAnnotator**: annotate points on arbitrary planes in 3D. This is useful for defining particle locations or other points of interest.
- **SphereAnnotator:** place spherical regions of interests in 3D volumes.
- **SurfaceAnnotator**: define a surface in an image. This is useful for labeling membranes.

### Supplementary Note 4: Manipulators

napari-threedee manipulators provide methods to interactively position objects in 3D. This is useful for refining positions and orientations of detected objects or exploring data by positioning rendering planes. We have developed 3 types of manipulators:

- **LayerManipulator**: Interactively position an image layer in 3D. This is useful for aligning multiple images (e.g., as a rough initial alignment for a registration algorithm).
- **PointManipulator:** Interactively position points in 3D. This is useful for positioning and orienting points in an image (e.g., picking particles in cryoelectron tomography).
- **RenderPlaneManipulator**: Interactively position a rendering plane in 3D. This is useful for viewing image volumes in non-axis-aligned planes.

### Supplementary Note 5: Visualizers

napari-threedee visualizers enhance how 3D data can be viewed in napari. This includes improved rendering and methods for exploring data. We have developed 3 types of visualizers:

- **CameraSpline:** Define a path along which you would like to explore your data using a spline annotator and then move the camera along that path. Like riding a roller coaster!
- **LightingControl:** Control the direction of lighting for a napari surface layer to shine light on important details.
- **AmbientOcclusion**: Apply shadows using ambient occlusion to a mesh layer to improve clarity of detailed features on the surface.

### Supplementary Note 6: documentation for napari-threedee

We have written in-depth documentation that is freely available online (https://napari-threedee.github.io/). This documentation is automatically built and served online via continuous integration (github actions). We have written tutorials and how-to articles to teach analysts how to use napari-threedee. Additionally, we provide an examples gallery to help users get started. For developers, we provide explanatory documentations for key concepts and API documentation describing the functions available in the library. We explain each type of documentation in more detail below.

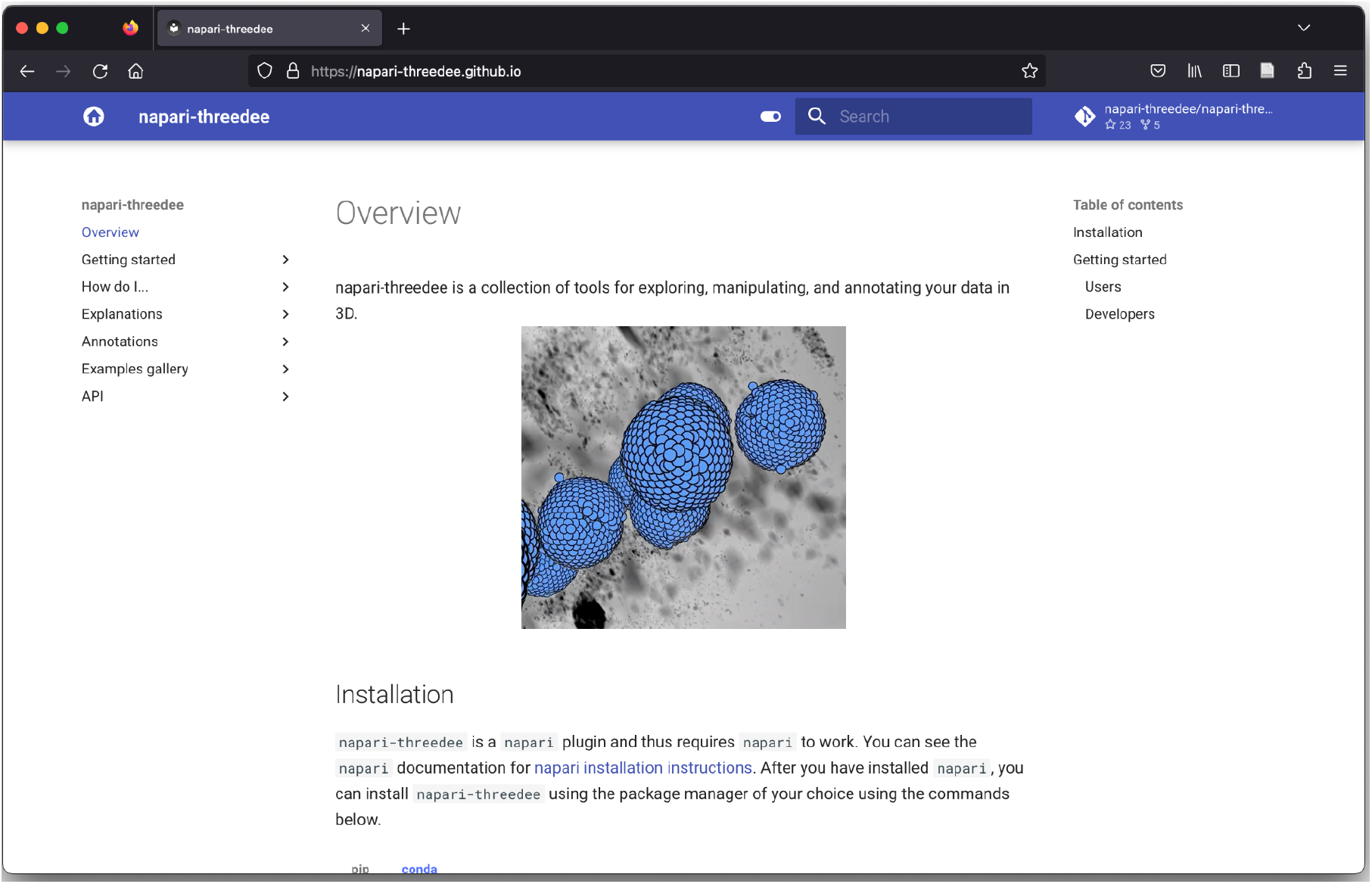

#### Tutorials and How-to articles

Tutorials provide end-to-end instructions on workflows or component usage. In particular, we have created a tutorial on how to use napari-threedee to label an image in 3D for a GPU accelerated segmentation workflow.

How-to documentation provides step-by-step instructions for specific operations. Thus far, we have provided 8 “how-to” articles . The “how-to” documentation can be found under the “How do I…” tab of the navigation bar.

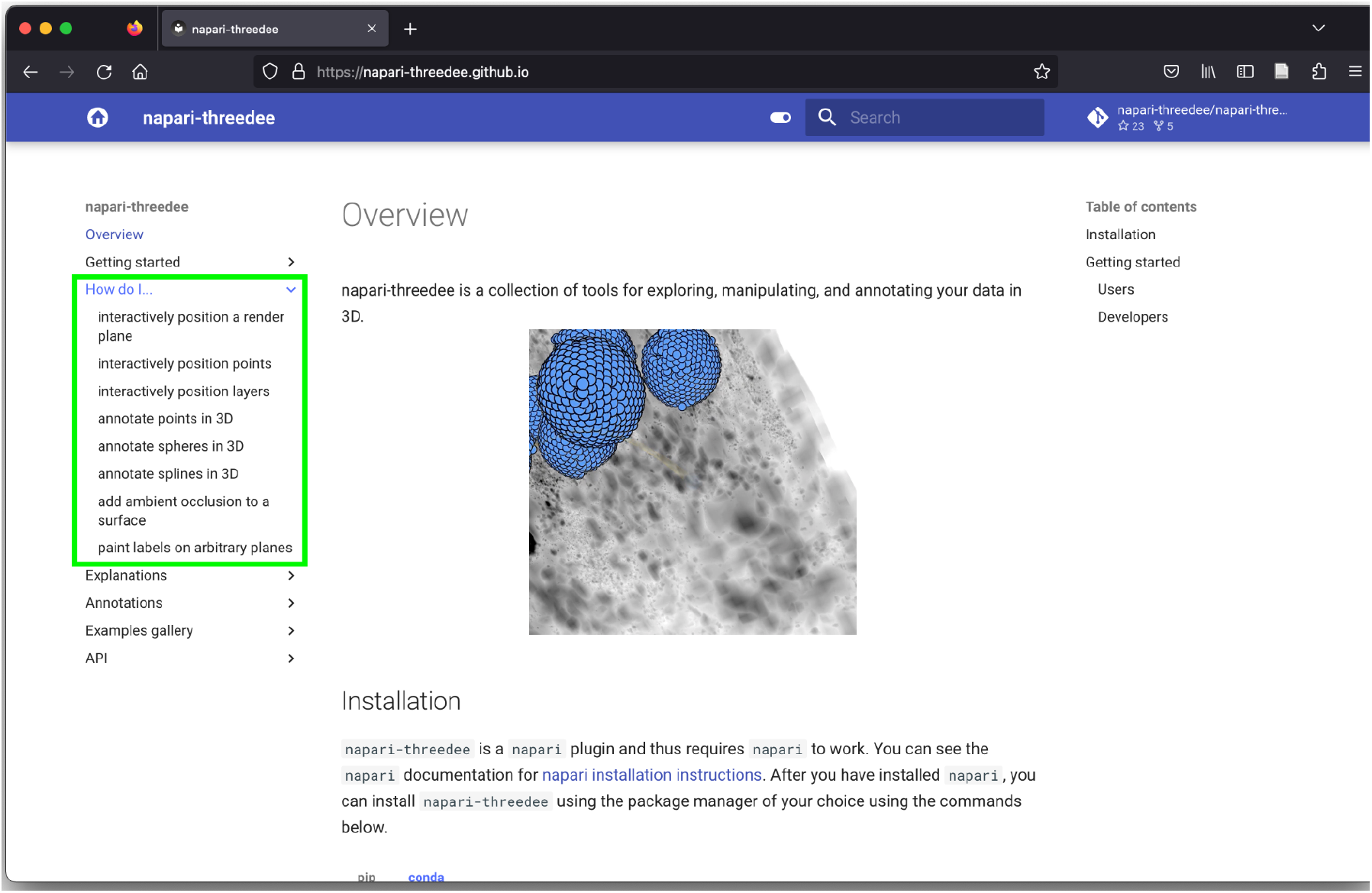

#### Examples gallery

To help users get started with napari-threedee, we have provided a gallery of example scripts that cover the usage of all napari-threedee components. Thus far, there are 20 example scripts available in the gallery. The gallery is automatically generated via github actions so it is kept up to date with the library. The gallery can be accessed here: https://napari-threedee.github.io/generated/gallery/

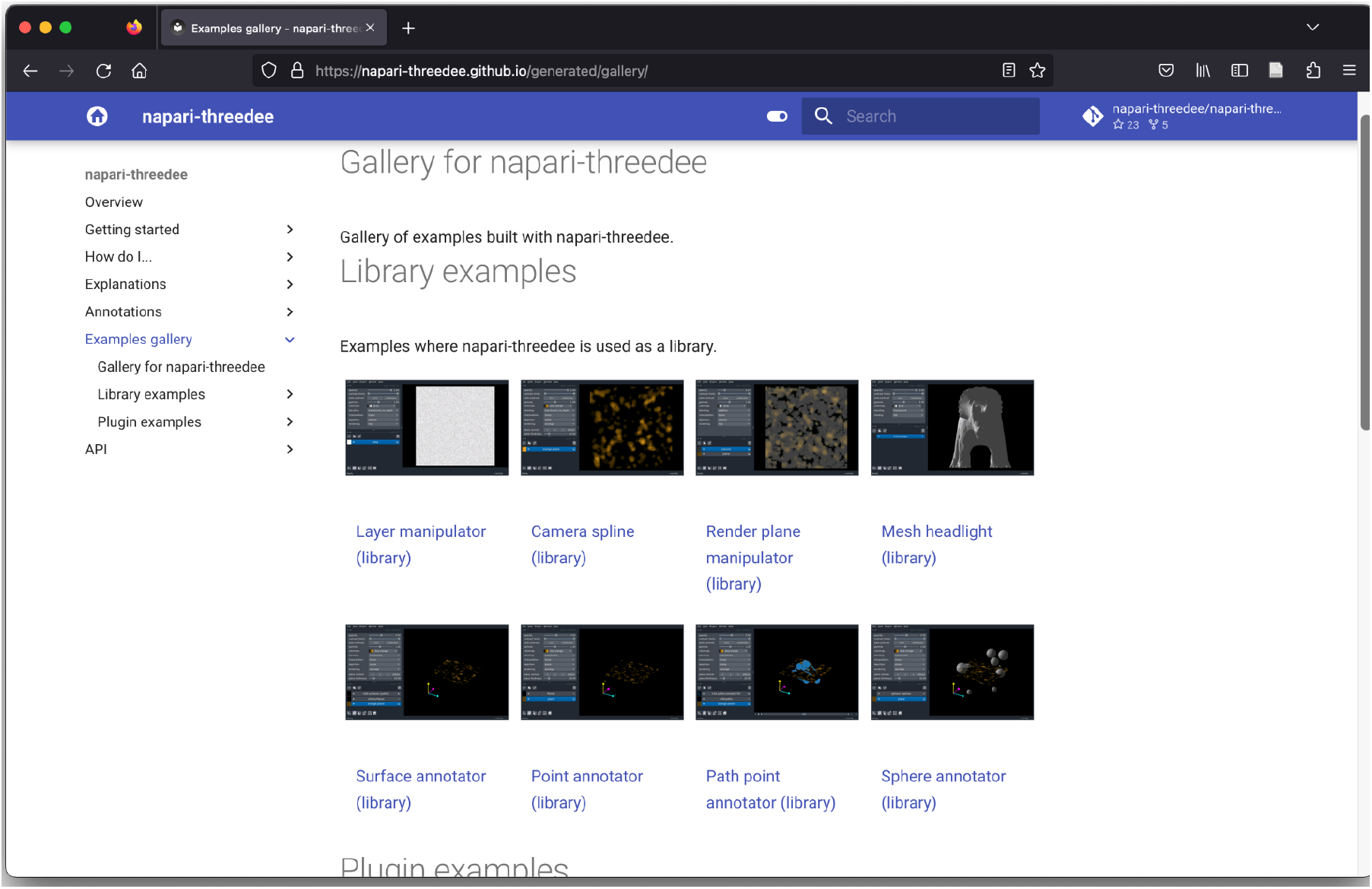

#### Explanations

Explanations are primers on technical concepts in napari-threedee. The intended audience is users and developers who are interested in learning how napari-threedee works. We have created explanatory documentation on:

| Topic | Link |
| --- | --- |
| How 3D interactivity works in napari | https://napari.org/stable/guides/3D_interactivity.html |
| Core concepts in napari-threedee | https://napari-threedee.github.io/dev_guides/core_concepts.html |
| How manipulators are designed | https://napari-threedee.github.io/dev_guides/manipulators.html |

#### API documentation

To help developers understand the napari-threedee codebase, we have automatically generated documentation describing the functions provided by the library.

